# Novel husbandry practices result in rapid rates of growth and sexual maturation without impacting adult behavior in the blind Mexican cavefish

**DOI:** 10.1101/2022.10.19.512864

**Authors:** Robert A. Kozol, Anders Yuiska, Ji Heon Han, Bernadeth Tolentino, Arthur Lopatto, Peter Lewis, Alexandra Paz, Alex C. Keene, Johanna E. Kowalko, Erik R Duboué

## Abstract

The development of animal model systems is dependent on the standardization of husbandry protocols that increase fecundity and reduce generation time. The blind Mexican tetra, *Astyanax mexicanus*, is an emerging genetic vertebrate model for evolution and biomedical research. Surface and cave populations of *A. mexicanus* have independently evolved, providing a model system for studying the genetic basis of divergent biological traits. While a rapid increase in the use of *A. mexicanus* has led to the generation of genetic tools including gene-editing and transgenesis, a slow and inconsistent growth rate remains a major limitation to the expanded application of *A. mexicanus*. The optimization of husbandry protocols that maximizes high-nutrient feed, smaller tank densities and larger tank sizes across development, would facilitate faster growth and expand the use of this model. Here, we describe standardized husbandry practices that optimize growth through a high protein diet, increased feeding, growth sorting of larvae and juveniles, and tank size transitions based on standard length. These changes to husbandry had a significant effect on growth rates and decreased the age of sexual maturity in comparison to our previous protocols. To determine whether our nutritional change and increased feeding impacted behavior, we tested fish in exploration and schooling assays. We found that a change in diet had no effect on the behaviors we tested, suggesting that increased feeding and rapid growth will not impact the natural variation in behavioral traits. Taken together, this standardized husbandry protocol will accelerate the development of *A. mexicanus* as a genetic model.

## Introduction

Husbandry for animal models has improved greatly over the last century, due to the scientific community’s desire to keep healthy stocks that create reproducible data ^1–3^. Over time changes in the size of tanks and housing for laboratory animals ^4–6^, improved diets ^7–9^ and environmental enrichment ^6, 10–12^ has provided healthier breeding stocks, while also boosting growth rates and fecundity. The majority of genetic and biomedical research is performed in a small number of models including *C. elegans*, *Drosophila*, zebrafish, and mice. A recognized commonality between these models is ease of husbandry, relatively fast generation time, and high fecundity ^1, 4–5, 8, 10^. These traits, along with optimized husbandry protocols have facilitated major discoveries and wide-spread use of these models.

As genome sequencing, along with mutagenesis and transgene technologies have become less expensive, non-traditional laboratory organisms are starting to be adopted by small communities ^14–18^. Furthermore, the expansion of genetic tools including genetic-editing have allowed for the expanded use of models to address diverse biological questions. Increasingly, these new models are being used to address the evolution of complex traits that are largely intractable in classic genetic models. Many of these models are challenging to rear in the lab due to slow generation time and low fecundity, presenting a major impediment to their use.

Aquatic organisms such as teleost fish provide advantages for studying the biology of vertebrates ^15, 19–22^. Many fish mate via dispersal spawning, releasing eggs and sperm into the water column at high numbers, which results in large clutch sizes that are easy to collect and maintain in small Petri dishes and bowls. Additionally, many embryonic and larval fish are transparent, which allows researchers to study internal organs, without euthanizing, dissecting or disturbing tissue ^17, 20, 23^. The blind Mexican tetra (*A. mexicanus*) is a teleost species that is emerging in the fields of evolution, development, and neuroscience ^24, 25^. The species is found in two distinct forms: an eyed, surface-dwelling form, and several hydrologically isolated populations of cavefish. Cavefish populations are the result of ancestral surface fish being washed into caves between 100,000-300,000 years ago ^26^, resulting in the convergence of traits, such as degenerated eyes ^27, 28^ and loss of pigment ^29, 30^, or loss of sleep ^19, 31^ and decreased stress ^32^. Although cavefish have been isolated and evolved troglomorphic phenotypes, surface to cave hybrids are viable and can be crossed for allele segregation and subsequent quantitative trait loci mapping ^25, 33, 34^.

Fish husbandry in the *A. mexicanus* community continues to vary across fish facilities and research groups. The recent implementation of gene editing^35, 36^ and tol-2 transgenesis ^14, 37^, along with the consistent generation of surface to cave hybrid populations, would benefit from shorter generation times, without jeopardizing health outcomes for reared fish. While our initial goal was to standardize feeding, we utilized variables tested in previous animal models ^4, 7–9, 38^, such as a high-nutrient diet, lower tank densities and feeding/tank size scaling with body length, to optimize growth rates and maximize welfare.

To define a standard protocol for optimizing growth rates, we used bimonthly observational and management periods to determine whether changes in diet, feeding frequency and tank conditions across growth improved rearing times. Our new husbandry protocol implemented several changes that includes; scaling food particle size to match caloric density to fish size, reducing tank densities (fish per liter) and sorting fish according to standard length (SL) to match food type and tank size. In our experience, these changes in husbandry resulted in temporal reduction to reach sexual maturity, from 8-10 months to 5 months post-fertilization. Finally, because feeding can have an impact on animal behavior, we wanted to test whether an increase in feeding frequency altered preexisting behavioral phenotypes for surface and cave populations. Fish raised in our growth rate optimized protocol show no difference in individual and group behavior. These results provide a clear rationale for adopting this protocol for *A. mexicanus* husbandry; high growth rates, lower mortality, and shorter generation time, with no observable impact on natural behavioral variation.

## Results

### Scaling nutrient density of food source with standard length provides high growth rates and fast generation times in both populations

Larval nutrient uptake is limited by the size of the mouth and digestive track during larval and juvenile periods, limiting growth during the first few months of development ^7, 39^. To provide a standardized diet, we increased feeding frequency and food particle size of high protein content feed across life stages, decreasing feeding effort while maintaining high nutrient value (Table 1). Larvae were initially fed freshly hatched artemia shrimp following 6 days post-fertilization (dpf) (Figure 1, Table 1). We monitored the growth of each fish across development, and by the time fish had a length equal to 10 mm (4-6 weeks), 300 μm Gemma (Figure 1b, Table 1) was added once a day as a supplement to hatched artemia. Following 6 weeks of development, larvae exceeding 12 mm standard length were further supplemented with blood worms segment tips (Figure 1c, Table 1) and monitored to ensure no fry struggled with ingestion. Studies have shown fish prefer bloodworms in comparison to other feed, providing an environmental enrichment that promotes feeding and growth ^40, 41^. At 15 mm (8-10 weeks), feeding was changed from Gemma to ground Zeigler pellets twice daily (Figure 1d) and blood worms (Figure 1e, Table 1) once a day. Zeigler pellets provide an improvement in nutrient density and decreases feeding time, due to particle size (Gemma = 300 μM, Ground Zeigler = ~1 mm). At 20 mm SL, fish were switched to full Zeigler pellets (3mm diameter) and blood worms twice daily. Finally, at 30 mm SL, feeding was increased to morning, afternoon, and evening. This feeding schedule was then followed through the end of the study, along with breeding protocols that were performed every two weeks. Fertilization of healthy embryos occurred at 42-50 mm SL, with time to sexual maturity showing variation between tanks but not between surface and cave populations (5-6 months). These results provide a feeding schedule across development to rapidly achieve sexual maturation in both surface and cave populations.

**Table 1.** Feeding schedule matching standard length to feeding type and frequency.

| Fish size in standard length | Feed Types (frequency) |
| --- | --- |
| 6-10 mm | Artemia (once) |
| > 10 mm | Artemia (once), Gemma 300 μm (once) |
| > 12 mm | Gemma 300 μm (once), blood worm segment tips (once) |
| > 15 mm | Ground Zeigler pellets (twice), blood worm segment tips (once) |
| > 20 mm | Zeigler pellets (twice), blood worms (twice) |
| > 30 mm | Zeigler pellets (thrice), blood worms (thrice) |

standard length = length of a straight line from snout to the end of the caudal peduncle

**Figure 1.**
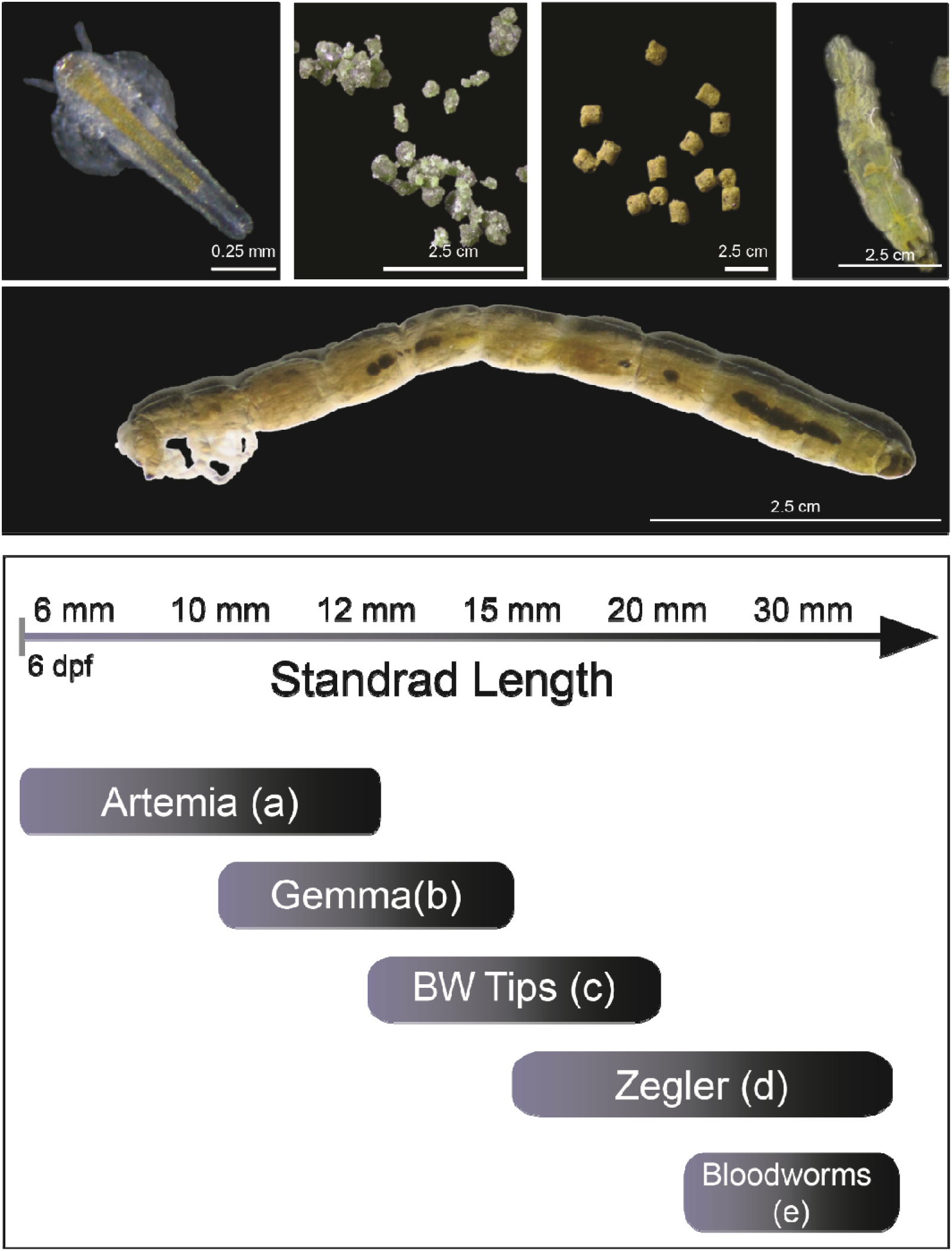
Food type and feeding schedule for high growth rate protocol. a. 24-hour hatched artemia nauplius. b. Gemma pellet feed. c. Zeigler pellets. d. Blood worm tips. e. Full blood worm. Arrow denotes fish size in standard length (mm; distance from nose to caudal peduncle).

### Sorting fish by standard length and creating a scalable tank size schedule results in low variation of growth across populations

Tank densities (fish per liter) have been shown to impact the growth and health of developing fish ^42 43, 44^. To ensure improved growth rates and optimal health outcomes, we created a schedule for increasing tank size based on standard length. Standard length was measured at bimonthly time points, beginning at 2 weeks of development, before the fish were put onto the aquarium system (Table 2&3, Figure 2). Cavefish exhibited a larger standard length in comparison to surface fish (Table 2), from four weeks to six weeks (Figure 2, Supplemental Table S1&2). Fish were then progressively sorted and placed in larger tanks (Table 3; 9 L & 5 Gal), until 40 mm standard length, when fish were finally moved to 10-gallon tanks. Fish exhibited exponential growth throughout the tank schedule changes, from Week 8 to Week 14, with average standard length increasing nearly 4-fold over those 8 weeks (Surface= 12.3 ± 0.6 mm to 40.4 ± 0.5 mm; Pachòn = 15.3 ± 0.5 mm to 42 ± 0.4 mm; Table 2, Figure 2). These results provide a standard for high growth rates through the coupling of tank size and standard length.

**Table 2.** Standard Length (standard length) in mm across 20 weeks of development.

|  |  |  |  |  |  |  |  |  |  |  |
| --- | --- | --- | --- | --- | --- | --- | --- | --- | --- | --- |
|  | 2 | 4 | 6 | 8 | 10 | 12 | 14 | 16 | 18 | 20 |
| SF | 7.5<br>± 0.1 | 12.6<br>± 0.2 | 13.5<br>± 0.6 | 19.7<br>± 0.4 | 26.3<br>± 0.6 | 34.1<br>± 1.5 | 41.0<br>± 0.5 | 43.3<br>± 0.3 | 45.1<br>± 0.2 | 48.1<br>± 0.2 |
| C | 8.0<br>± 0.2 | 13.3<br>± 0.4 | 15.3<br>± 0.5 | 24.3<br>± 0.7 | 26.9<br>± 0.6 | 34.0<br>± 0.9 | 41.6<br>± 0.4 | 46.1<br>± 0.4 | 47.0<br>± 0.3 | 50.2<br>± 0.2 |

± = standard error.

**Table 3.**
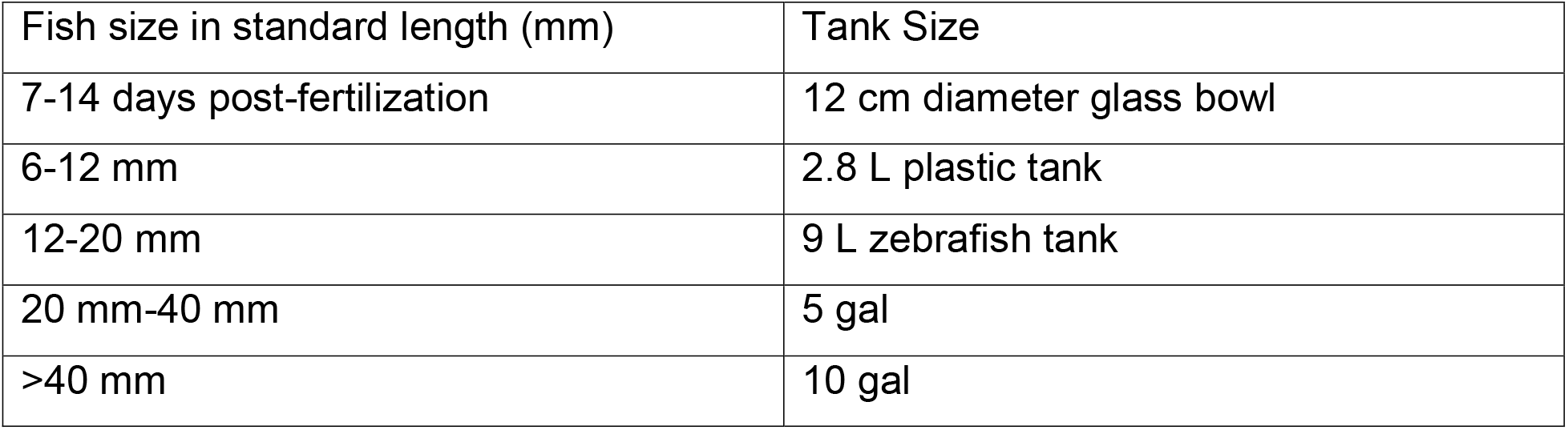
Schedule for changing tank sizes across development.

| Fish size in standard length (mm) | Tank Size |
| --- | --- |
| 7-14 days post-fertilization | 12 cm diameter glass bowl |
| 6-12 mm | 2.8 L plastic tank |
| 12-20 mm | 9 L zebrafish tank |
| 20 mm-40 mm | 5 gal |
| >40 mm | 10 gal |

**Figure 2.**
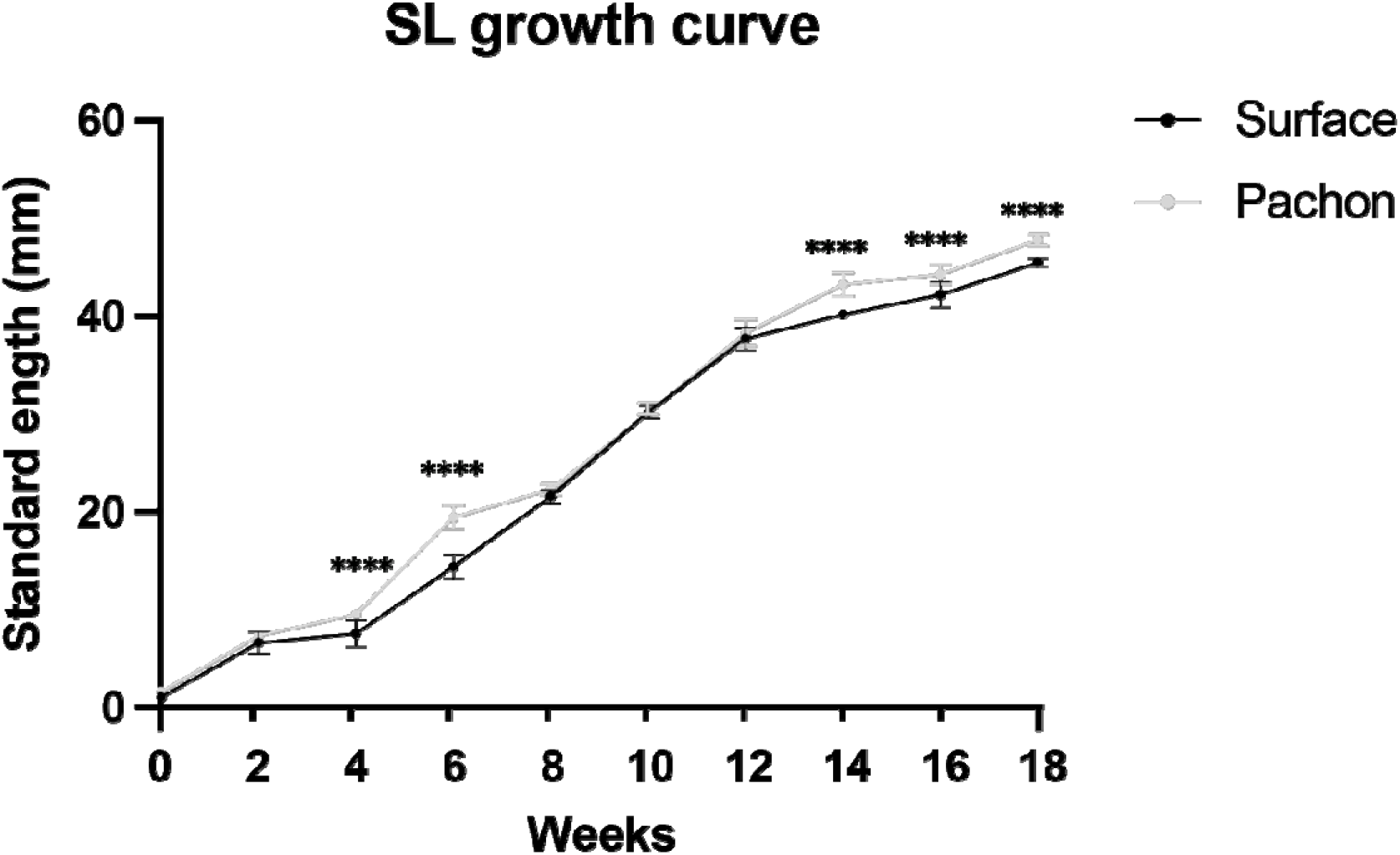
Standard length measurements of surface fish and Pachón cavefish through 5 months of development. Standard length measurements in millimeters (y-axis) was recoded every two weeks (x-axis) until fish reached 50 mm in standard length. X-axis starts at week 2 when fish were measured and put onto the recirculating system in the fish facility. Error bars denote +-standard error. Sample size, n=24 for both surface fish and cavefish. Unpaired t-test with significance values; * = p<0.05, ** = p<0.01, *** = p<0.001, **** = p<0.0001.

### High protein meals that are staged matched for feeding frequency reduces mortality rates and improves adult welfare

We monitored mortality and tank density throughout development to determine what stages contributed to mortality, and whether increasing the frequency of feeding impacted health and tank behavior. Larvae were added to the fish facility system at a density of 16 fish per tank (2.8L). Mortality rates for fry during the first month averaged 22% (± 0.02) across populations, resulting in a tank density of 10-13 fish per tank (average=12 ± 0.22 fish; Table 4). Following the first month of development (8-14 mm), we did not record further mortality in any tank for all populations. In addition to low mortality rates, we also observed few instances of surface fish exhibiting attacking behavior (reviewing video for growth recordings every two weeks over a five-month period). Furthermore, throughout the duration of the study, only one fish needed to be isolated due to aggression by siblings. Overall, our new feeding strategy, tank size and sorting scheduling results in low mortality rates and aggression.

**Table 4.** Tank population size and mortality rate.

|  | Tank 1 | Tank 2 | Tank 3 | Tank 4 | Tank 5 | Average |
| --- | --- | --- | --- | --- | --- | --- |
| SF | 13 (0.13) | 11 (0.26) | 12 (0.20) | 12 (0.20) | 11 (0.26) | 11.8 (0.22) |
| C | 12 (0.20) | 13 (0.13) | 10 (0.33) | 11 (0.26) | 12 (0.20) | 11.6 (0.23) |

(#) = mortality rate of tank,

### Exploration behavior and schooling in adult surface fish and cavefish populations are not impacted by changes in husbandry practices

Changes to animal care can impact animal behavior and therefore influence the reproducibility of behavioral studies^10, 45, 46^. To determine whether our changes in animal care could impact behavior, we utilized standard protocols for published behavioral assays at the individual and group level ^32, 47^. Behavioral and physiological indicators of stress are reduced in cavefish compared to surface fish ^32^. The novel tank assay measures stress response from determining the ratio of time spent in the top (low stress) versus bottom (high stress) ^32^. Our results faithfully recapitulate previously published results, with surface fish displaying high stress (bottom dwelling) and Pachòn cavefish exhibiting low stress (top dwelling) regardless of husbandry protocol (Figure 3., Supplemental Tables S3-6). Finally, to determine how group behavior is impacted by our new protocol, pairs of adult *A. mexicanus* were observed for schooling behavior in a 111 cm diameter tank for 20 minutes. Again, fish raised with our new standard protocol recapitulated previously published data ^47^, with Pachòn cavefish lacking coherent schooling behaviors, while surface fish display schooling behaviors (Figure 4, Supplemental Tables S7-10). These results suggest our changes to husbandry do not impact well-established individual and group behavior in surface fish and cavefish populations of *A. mexicanus*. Therefore, these protocols increase the growth and survival of this model without impacting complex behaviors commonly studied in this system.

**Figure 3.**
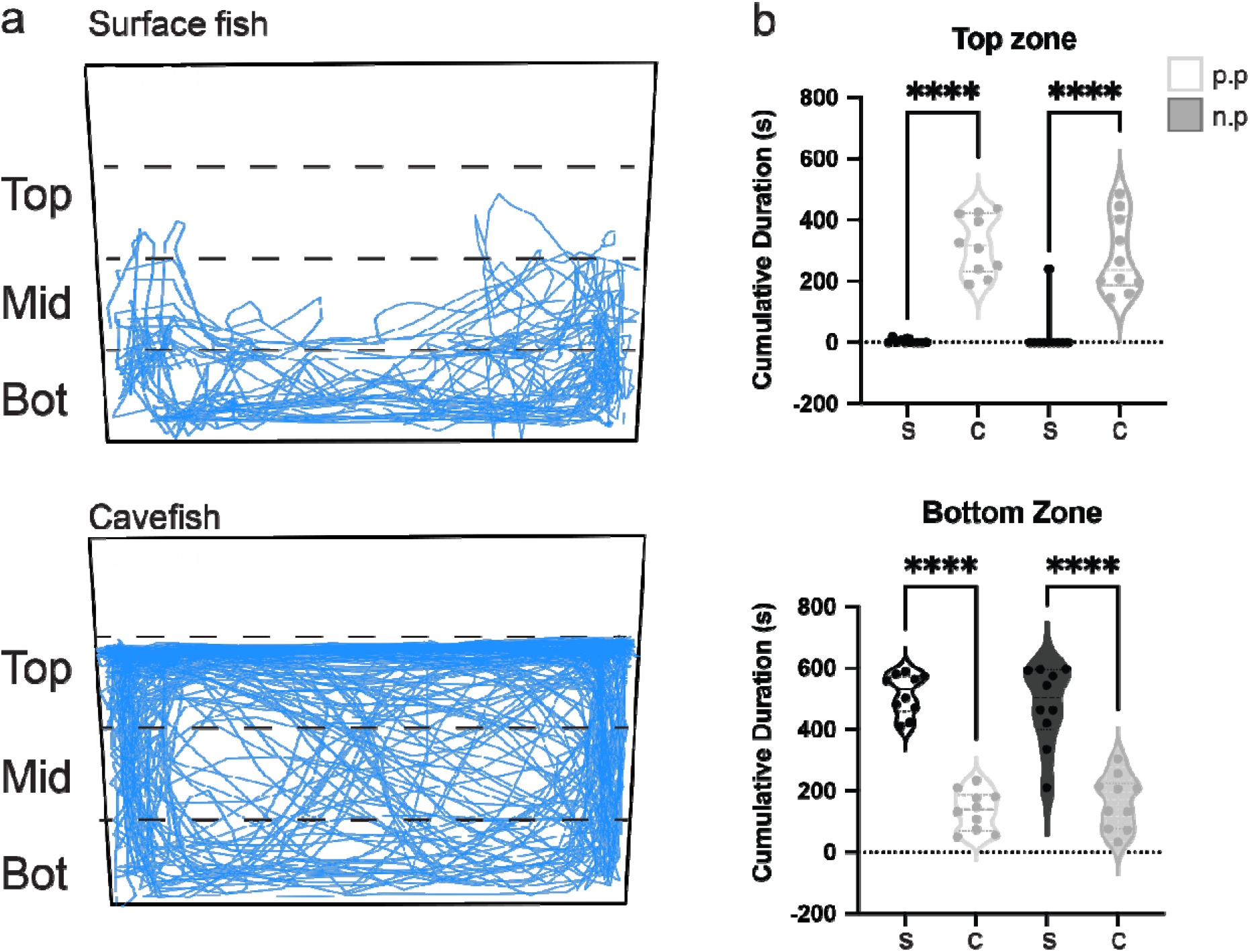
Novel tank assay comparing adult Astyanax populations raised with previous and high growth rate husbandry protocol. a. Tracking traces (blue) from individual novel tank trials, with grey shading representing water column. b. Cumulative duration of time spent in the top and bottom half of the tank. Previous husbandry protocol (p.p) and new husbandry protocol (n.p.). Violin plots display min to maximum values, with three lines representing the 75th quartile, median and 25^th^ quartile. Sample size (n = 10) were the same for all populations. P value asterisks; **** = p<0.0001.

**Figure 4.**
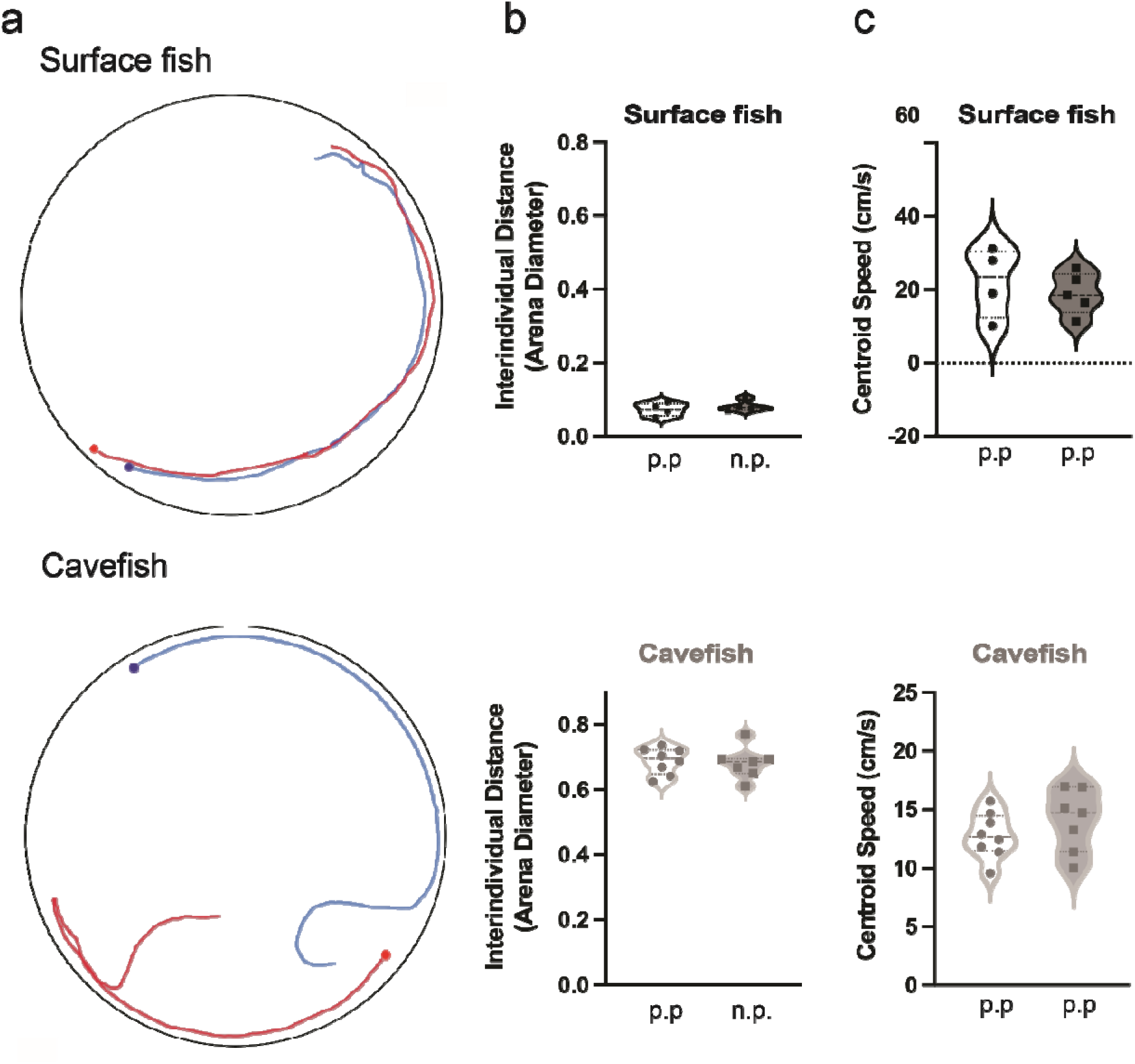
Schooling assay comparing adult Astyanax populations raised with previous and standardized husbandry. a. Swimming tracks representing a single trial of two fish from the same population (e.g., red =fish 1, blue = fish 2). b. Violin plots of interindividual distance, the average distance in cm between both fish in each trial. c. Violin plots of centroid speed, average speed calculated from tracking a centroid placed at the middle of each fish. Previous husbandry protocol (p.p) and new husbandry protocol (n.p.). Violin plots display min to maximum values, with three lines representing the 75th quartile, median and 25^th^ quartile. Sample sizes for surface fish, standard n = 4 and optimized n = 5. Sample sizes for cavefish fish, standard n = 8 and optimized n = 7. No comparisons were statistically significant.

## Discussion

A standardized husbandry protocol is necessary as the cavefish community grows across the globe. Cavefish provides a model with high-genetic diversity for direct genotype-phenotype research in relation to evolution and disease. However, the low growth rates and larger tank sizes needed for cavefish husbandry in comparison to other aquatic models ^48^ make development of new generations more costly in terms of time and fish capacity. Here we show that a progression in particle size of high nutrient feed, tank density, tank size and uniform standard length sorting provides a high growth rate that can result in healthy breeding adults by 5 months post fertilization. This protocol will greatly decrease the time it takes to develop hybrids lines, along with stable transgenic and CRISPR mutated fish. With the growth and health outcomes achieved, coupled with experiments showing that a change in diet did not affect behavior, we suggest that cavefish community members improve their adult generation timetable by adopting this husbandry protocol.

### Enriched early life diet and increased feeding frequency resulted in accelerated growth

Past laboratory studies have commented on food sources for *A. mexicanus* but vary in the types of feed and feeding frequencies ^14, 49^. We previously fed populations of *A. mexicanus* artemia for larval and juvenile periods, switching to flake food twice daily for adults. Therefore, we decided to utilize a typical zebrafish feeding regime, moving up in particle size as the fry grow, scaling appropriately with jaw and stomach size ^38^. In this study, we went from artemia, to small grain Gemma, before scaling up to Zeigler pellets and blood worms. We also decided to omit flake food that loses its nutrient value while sitting in the water column ^50^. In the zebrafish community, 2-3 feedings per day is recommended because fish lack a true stomach and therefore pass through quickly allowing for increased feeding ^51, 52^. Our goal was to raise 50 mm standard length adults that could produce healthy embryos. This goal was achieved with a clear exponential growth period (week 6 to week 18) that may provide a critical period of development for reaching a larger size, along with sexual maturity in a reasonable timetable.

### Changes in diet, tank density and tank size for standard length improved mortality rates and appears to lower aggression in surface fish tanks

Importantly, we observed a low mortality rate during development and overall decreased aggression in adult surface fish populations. While some mortality is inevitable, due to growth defects and competition for food, we found that the majority of mortality occurs around the end of the first month, before fish display exponential growth. Furthermore, a survival rate of 78% mirrors high survival rates observed in updated zebrafish husbandry methods ^38, 53^. Also in the current study, we did not record a single mortality following 6 weeks of development. We find an average tank density of 2-3 fish per gallon, results in steady growth and good health outcomes for developing fish. Additionally, smaller fish can be bullied in adult populations, that ultimately results in bodily injury to bullied fish and at worst cases death ^54, 55^. Surprisingly, we only observed one incident that resulted in the transfer of the bullied fish for recuperation. Lack of fighting/biting injuries was later confirmed by reviewing images and videos taken for length measurements, revealing no observable injuries to surface fish across growth periods. These results suggest that, not only is our revised protocol ideal for quickly raising adults, but it improves community health by lowering aggression.

### No observed behavioral change in fish raised with optimized protocol

Several zebrafish studies have shown that changes in diet and environment can affect a fish’s physiology and behavior ^45, 56, 57^. Due to the importance of reproducibility in scientific research, it is paramount that biological data is consistent across labs and does not vary due to changes in diet and husbandry practices. We found no change in adult exploration or schooling behavior of either surface fish or cavefish when comparing standard to optimized protocol reared fish. One recent zebrafish study showed lower feeding frequency increased anxiety in adult zebrafish, which may help explain the reduction in bullying that we observed with more feeding ^57^. Certainly, an increase in tank size, density, and an additional feeding period per day, could provide enough resources across 24 hours to reduce competition. Overall, this study provides a standardized way to increase growth rates and decrease time to sexual maturity, while still maintaining population specific phenotypes that are necessary for studying the evolution of complex traits.

## Conclusion

Our results provide a husbandry protocol that facilitates the enhanced use of *A. mexicanus*. This is likely to greatly improve existing applications and experiments, such as adult fish research, replenishment of breeding stocks, and generating hundreds of hybrid fish necessary for QTL mapping. Our findings clearly show that a change in diet and management of populations across growth periods can drastically reduce the time it takes to raise fish to adulthood, while also improving mortality and maintaining a healthy behavioral environment.

## Materials and methods

### Fish maintenance and husbandry

*A. mexicanus* were cared for in accordance with NIH guidelines and all experiments were approved by the Florida Atlantic University Institutional Care and Use Committee protocol #A1929. *A. mexicanus* stocks were housed in the Florida Atlantic Universities Mexican tetra core facilities. *A. mexicanus* fish lines used for this study; Pachón cavefish stocks were initially derived from Richard Borowsky (NYU); Surface fish stocks were derived from Rio Choy wild populations.

### Dietary ingredients and feeding schedule

Control fish stocks were fed using pet store flake food with a nutritional content of 40% crude protein, 5% crude fat, 5% crude fiber, 9% moisture, and 9% ash. Fish fed under our new protocol changed diet during development; Gemma as larvae, ground blood worms and Zeigler pellets as juveniles, and blood worms and Zeigler pellets as adults. Brine shrimp – 60% protein, 24% fat, 4.4% ash and 8.5% moisture - was used as a supplement at all stages. Gemma (100-500 pellet size) nutritional content; 59% protein, 14% oil, 14% ash, 0.2% fiber and 1.3% phosphorus. Zeigler pellets nutritional content; 45% protein, 16% fat, 2% fiber, 12% moisture, and 8% ash. Fish were fed to satiation and tank cleaning was performed every other week to promote excellent water quality and high oxygenation.

### Standard length measurements

Videos were collected with rulers placed at the front and back of the tank. Frames were analyzed in Fiji by measuring standard length of fish swimming against the ruler’s edge. Scale was coded by setting the pixel length of 1 cm line segment against the ruler. Five fish per tank were measured for each time point and continued measuring through 5 cm in standard length. A standard length of 50 mm or 5 cm was chosen as a target because previous laboratory members have reported successful breeding of 4-5 cm standard length surface fish and cavefish (data not shown).

### Novel Tank Assay

Adult fish were transported to the adult behavioral room to acclimate for 1 hour. Following acclimation, individual fish were added singly to 5L plastic zebrafish tanks (Aquaneering Inc., San Diego, CA, USA) and recorded using a FLIR Grasshopper^®^3, GS3-U3-23S6M-C 1/1.2 (Teledyne FLIR LLC., Wilsonville, OR, USA) at 15 frames per second for 15 minutes. Videos were then analyzed using Noldus EthoVision^®^ Software XT14 (Noldus Inc., Wageningen, Netherlands, EU) to track and measure time spent at the top and bottom half of the tank.

### Schooling Assay

Fish were fed at least 1 hour prior to being assayed and then carried to a designated behavior room in a 2.5 L carrier tank. Pairs of fish were then gently netted into the experimental arena and allowed to acclimate for 10 minutes. All individuals that were assayed together were from the same home tank. Experiments were conducted in a round tank (111 cm diameter × 66 cm height) filled to a depth of 9 cm with system water. A Genius WideCam F100 video camera (Dongguan Gaoying Computer Products Co., Guangdong, China) was affixed to a custom-built PVC stand that allowed recording from above the center of the tank. Lighting was provided via four white 75-watt equivalent halogen light bulbs (Philips A19 Long Life Light Bulb, Amsterdam, Netherlands) mounted in clamp lights with 5.5 in shades (HDX, The Home Depot, Georgia, United States) to diffuse light. Videos were collected at 30 fps using OBS Studio (Open Broadcaster Software).

Automated tracking was done using EthoVision XT v. 13.0.1220 and data were exported as .xlsx files containing the X and Y coordinates of each fish. The median pair distance between and median centroid speed were found for each trial using the Pandas and NumPy libraries in Python 3.9.5. Pair distance was calculated as the distance between the center points of each fish and centroid speed was calculated as the movement speed of the center point directly between individuals. The Shapiro-Wilk test was performed to assess normality of pair distance and centroid speed medians. Data were subsequently compared using unpaired t-tests. All statistical analyses were performed using GraphPad Prism 9.2.1.

## Supporting information

Statistical Tables

## Acknowledgements

This work was supported by NSF EDGE grant 1923372 (ERD and JEK), NSF grant IOS2202359 (JEK), NIH grant R35GM138345 (JEK), and NIH grant R24OD030214 (ACK).

## Notes

https://drive.google.com/drive/folders/1rmdiwb_CWrFbDTVMBvFWddrhdBMbN-qf?usp=sharing

### Competing Interest Statement

The authors have declared no competing interest.

